# Can CRISPR-based gene drive be confined in the wild? A question for molecular and population biology

**DOI:** 10.1101/173914

**Authors:** John M. Marshall, Omar S. Akbari

## Abstract

The recent discovery of CRISPR and its application as a gene editing tool has enabled a range of gene drive systems to be engineered with much greater ease. In order for the benefits of this technology to be realized, drive systems must be developed that are capable of both spreading into populations to achieve their desired impact, and being recalled in the event of unwanted consequences or public disfavor. We review the performance of three broad categories of drive systems at achieving these goals - threshold-dependent drives, homing-based drive and remediation systems, and temporally self-limiting systems such as daisy-chain drives.

## Introduction

The recent discovery of CRISPR and its application as a gene editing tool has enabled gene drive systems to be engineered with much greater ease (*1, 2*). Recent attention has been focused on homing-based gene drive systems; however the ease of gene editing has advanced the entire field of gene drive, enabling scientists to implement a wide range of drive architectures, many inspired by naturally-occurring systems found in the wild (*2, 3*). Gene drive technologies have the potential to provide revolutionary solutions to a broad range of public health and environmental issues - for instance, controlling insect-borne diseases, removing invasive species, and reversing the development of resistance to insecticides and herbicides, all in an economically viable and environmentally friendly manner. Advantageous traits could also be spread more quickly into populations, compared to the case of natural selection, providing a potential solution to preserve endangered species threatened by pathogens.

In order for these benefits to be realized, gene drive systems must be developed that are both: a) capable of effectively spreading into populations to achieve the desired agricultural or epidemiological impact, and b) able to be recalled from the environment in the event of unwanted consequences or public disfavor. With this in mind, an ideal gene drive system would be capable of spreading to fixation or near-fixation locally; but without spreading on a wide scale, so that efforts to remediate it from the environment may be confined to a manageable area. As the technology matures and implementation becomes within reach, the question of the extent to which gene drive can be limited in the wild gains operational relevance.

Here, we address this question as it applies to three broad categories of drive systems, all of which can likely be engineered more easily using CRISPR: a) threshold-dependent drives, b) threshold-independent drives, and c) temporally self-limiting drives. Threshold-dependent drives are distinct in the sense that they must exceed a critical frequency in the population in order to spread (*4, 5*). These systems are well-suited for local confinement and may be eliminated from a population through being diluted below the threshold frequency. Threshold-independent systems, on the other hand, are able to spread from very low initial population frequencies (*6*–*10*). These systems are at higher risk of spreading into neighboring populations and countermeasures have been proposed to remediate them from the environment in order to limit their spread (*1, 2, 11*–*13*). Thirdly, temporally self-limiting systems display transient drive activity and may disperse somewhat into neighboring populations, but are eventually eliminated by virtue of fitness costs (*14, 15*). For each of these systems, we review their molecular and population biology, and discuss whether the molecular tools currently being developed are capable of meeting the challenges of confining drive in the wild.

## Threshold-dependent drive systems

### Translocations

The first gene drive system proposed to control insect pest species was a threshold-dependent one - reciprocal chromosomal translocations (*16*). In a landmark paper, Curtis (1968) showed that if a chromosomal translocation was inexorably linked to a desirable gene such as one conferring malaria refractoriness in a mosquito, then it could spread that gene into a population provided it were released above a critical threshold frequency. In the 1970’s-80’s, several attempts were made to generate reciprocal chromosomal translocations in mosquitoes to control wild populations; however these efforts were unsuccessful and the approach was ultimately abandoned (*17, 18*). New efforts have refounded hope for this technique as precise chromosomal translocations were recently engineered using endonucleases and shown to drive in laboratory experiments with a threshold frequency of ~50% (Figure 1A) (*5*). Site-specific chromosomal translocations have recently been engineered using CRISPR in several species, suggesting it is only a matter of time until CRIPSR-based translocation drives will be widely available (*19*–*21*).

**Figure 1.**
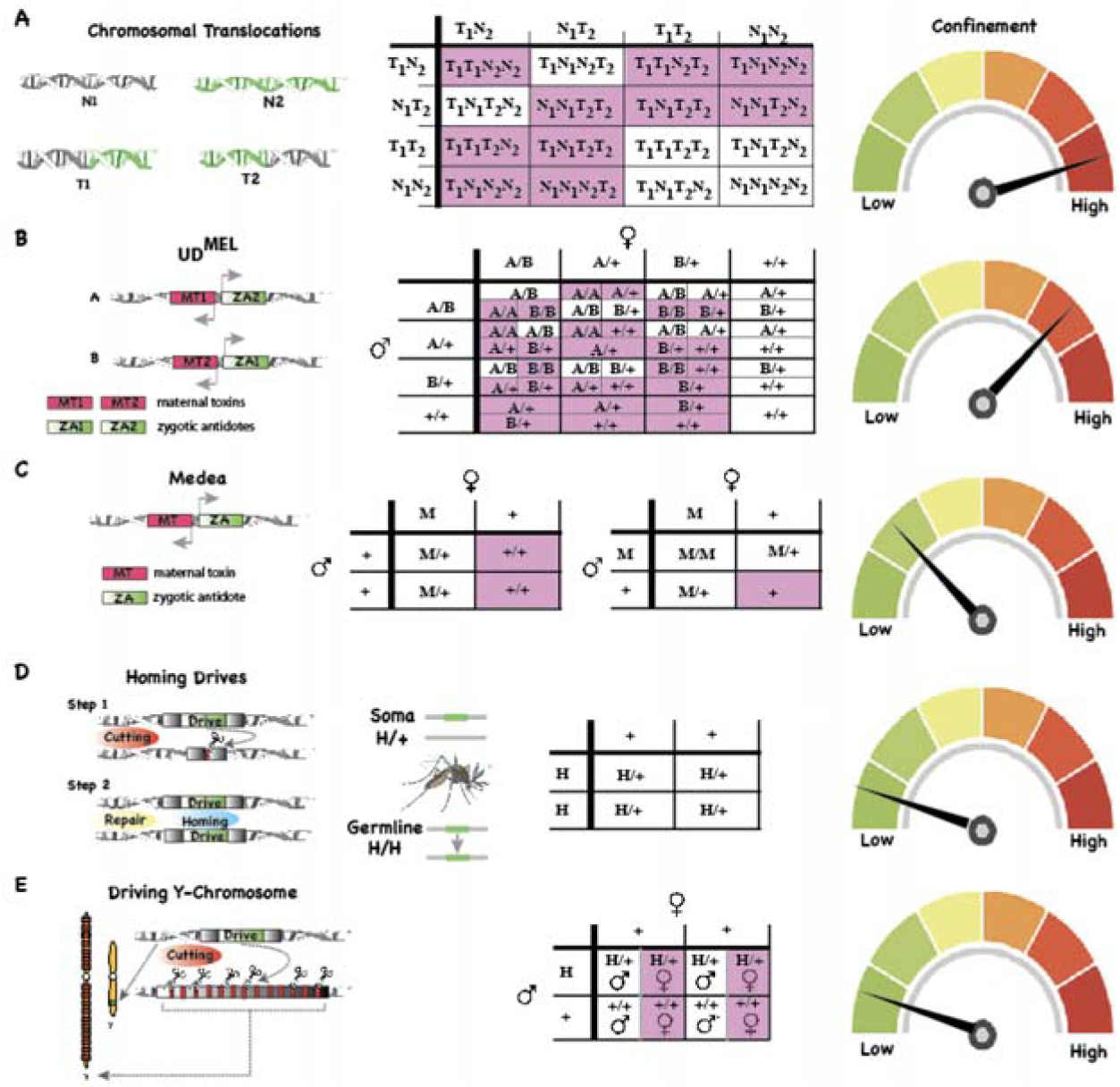
Various classes of gene drive systems illustrating mechanism, inheritance patterns and confinement properties. Reciprocal translocations (T1 and T2) result from the mutual exchange between terminal segments of two non-homologous chromosomes (N1 and N2). When translocation heterozygotes mate with each other, there are several patterns of disjunction resulting from independent meiotic assortment. Given that half of the gametes are unbalanced and many combinations of gametes produce unviable offspring, translocations display threshold-dependent dynamics and are expected to be confinable to local populations (A). UD^MEL^ is composed of two constructs each consisting of a zygotically-expressed antidote and a maternally-expressed toxin. Each toxin (MT1 or MT2) is linked with an antidote (ZA1 or ZA2) for the opposing toxin. As a result, the offspring’s chances of inheriting the proper combination is frequency-dependent and represents a form of underdominance that is expected to be confinable to local populations (B). *Medea* systems rely on the timed expression of a maternal toxin (MT) and zygotic antidote (ZA). This combination results in the death of all progeny that fail to inherit the *Medea* allele from a heterozygous mother, resulting in drive from low initial population frequencies (C). Homing-based drives encode endonuclease machinery in the germline allowing them to cleave their competing allele and copy themselves. This results in most progeny of heterozygotes receiving the drive element enabling rapid spread of the homing system through a population (D). A synthetic driving Y chromosome encodes a nuclease, inserted on the Y chromosome, that cleaves repeated sequences exclusive to the X chromosome during spermatogenesis. This results in disruption of X-bearing sperm leading to predominantly Y-bearing sperm and therefore male offspring. This system is expected to rapidly increase in frequency in a population eventually leading to an all-male population crash (E).

### Engineered underdominant systems

Systems that manipulate inheritance by expressing toxins and antidotes at various stages of development can also display threshold dynamics (*22*). This can happen because, at low population frequencies, the toxin confers a significant fitness cost; but at high population frequencies, the antidote confers a selective benefit in the context of a prevalent toxin. A few systems displaying this property were proposed by Davis *et al.* (*23*), and recently the first synthetic, threshold-dependent gene drive system, relying on the expression of counteracting toxins and antidotes, was engineered in the laboratory (*4*). This underdominant system, called UD^MEL^, consists of two unlinked constructs, each possessing a maternally-expressed toxin active during oogenesis, and a zygotically-active antidote expressed by the opposite construct (Figure 1B). The resulting dynamic, confirmed in laboratory drive experiments, is gene drive for releases frequencies above ~20%. To date this system has only been developed in a model organism, *Drosophila melanogaster*, primarily due to the difficulty of engineering miRNA toxins. However, the advent of CRISPR technology is expected to make it easier to engineer this system in other species as CRISPR gene editing may accelerate the development of maternal toxins (*1*).

### Population biology

Anticipating how these systems spread in the wild requires a detailed population modeling framework specific to the species of interest. Simple population models, in which two randomly-mating populations exchange small numbers of migrants with each other, predict that these systems can be released at high frequencies in one population and spread to near-fixation there, but never take off in the neighboring population(s) because they do not exceed the threshold frequency there (*22*). If this dynamic held true in the wild, it would be an ideal scenario for local gene drive: a) the system would spread at its release site following an intentional release, b) it would only spread into neighboring populations at very low levels, and c) it could be eliminated through dilution with wild-type organisms. However, whether this holds true or not depends crucially on the dispersal patterns and population structure of the species being considered. Species displaying moderate dispersal and infrequent inter-population movement, such as the malaria vector *Anopheles gambiae* (*24*), may be ideal candidates for such systems. Species displaying relatively less dispersal, such as the arboviral vector *Aedes aegypti* (*25*), may still be good candidates but may require more comprehensive releases to prevent the drive system from not reaching some population patches.

## Threshold-independent drive systems

### Medea

The first synthetic gene drive system was a threshold-independent one, *Medea* (Maternal Effect Dominant Embryonic Arrest), and displays a low-to-moderate threshold in the presence of a fitness cost. In a highly influential paper, Chen *et al.* showed that the *Medea* dynamics observed in natural systems (*26*), in which non-*Medea* offspring of *Medea* mothers are unviable, could be replicated synthetically through the action of a maternally-expressed toxin linked to a zygotically-expressed antidote (*10*). The antidote consists of a recoded version of the target gene that is immune to the effect of the toxin, and is expressed zygotically in embryos that inherit the *Medea* element (*9, 10, 27*). This potent toxin-antidote combination confers a selective advantage to the *Medea* construct, enabling it to spread into a population from very low initial frequencies (Figure 1C). *Medea* constructs engineered thus far function through the action of RNA interference-based toxin-antidote combinations in which synthetic miRNA toxins are expressed during oogenesis in *Medea*-bearing females, disrupting an essential embryonic gene in all progeny; however these have been difficult to engineer in some species, and it is hoped that CRISPR gene editing will accelerate the development of maternal toxins in the near future (*1*).

Like Curtis’ proposition for translocations, *Medea* has been envisaged as tool to drive desirable genes into populations; however, if the payload gene is no longer desired in the population, then mechanisms have been proposed to remove it. One proposition is to introduce a second-generation *Medea* element having a distinct maternal toxin to the first and having zygotic antidotes to both the first and second-generation toxins (*10*). Such a construct is expected to spread at the expense of both the first-generation *Medea* element and wild-type allele, with the goal of removing the effector gene from the population, albeit while leaving behind a residual *Medea* toxin-antidote machinery (Figure 2A). It should be noted that, for both drive and remediation to be effective, the toxin and antidote must be highly efficient. A naturally-occurring allele conferring resistance to the maternal toxin will be favored as drive is occurring, and in fact this was observed in a *Medea* element recently synthetically engineered in *D. suzukii (27).* Engineering efforts must therefore consider toxin-resistant alleles and how to engineer systems that are resilient to them.

**Figure 2.**
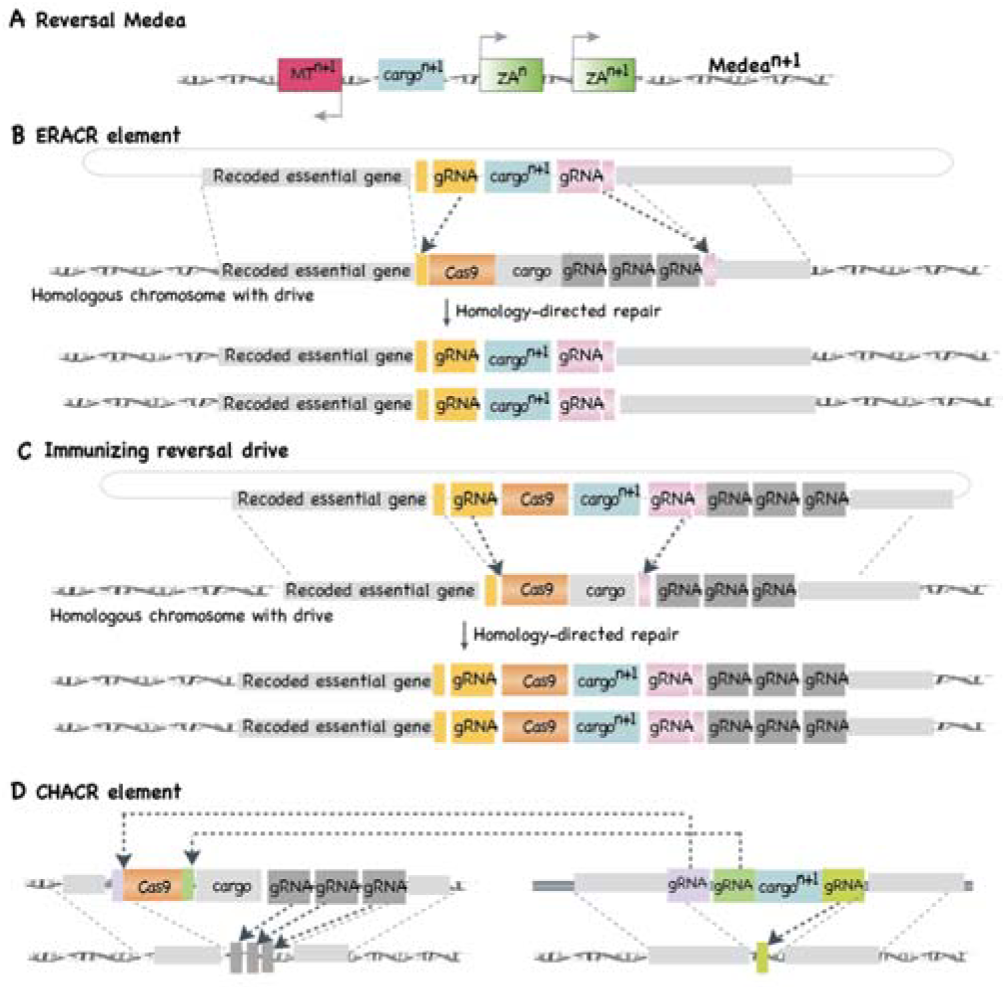
Remediation for threshold-independent drives. A first-generation *Medea* element can be remediated from the environment using a second-generation or reversal *Medea* element consisting of a new maternal toxin (MT^n+1^), a new effector gene (cargo^n+1^), the original zygotic antidote (ZA^n^), and a new zygotic antidote that is resistant to MT^n+1^ (ZA^n+1^) (A). An ERACR element can be used to remediate a fixed homing system by encoding guide RNAs (gRNAs), utilizing Cas9 from the original drive, that enable the ERACR to home into and inactivate the original drive (B). An immunizing reversal drive is similar to an ERACR, except that it encodes Cas9 and gRNAs that enable it to both remove the original drive system and to “immunize” wild-type individuals against the original homing system by homing into its corresponding target site (C). A CHACR encode several gRNAs that utilize the Cas9 from the original drive system that is to be removed. These gRNAs are designed to inactivate this original drive in addition to cleaving the allele opposite the CHACR to allow for copying of the CHACR via homing (D).

### Homing-based systems

The most quintessential form of drive, and one that can result in threshold-independence even in the presence of a fitness cost, occurs through the action of “homing.” First proposed by Austin Burt, homing-based drive systems use a sequence-specific endonuclease to cleave a highly-specific target sequence usually only present at a single site in the host genome (*28*). They are then copied to the opposite chromosome when the cell’s repair machinery uses this sequence as a template for homology-directed repair (Figure 1D). Early efforts to engineer these systems were implemented using homing endonuclease genes (*29*); however, the advent of CRISPR-Cas9 technology has greatly increased the ease of gene editing and homing-based gene drive systems are now straightforward to engineer (*1, 2*). In fact, in the last three years, CRISPR-Cas9-based homing drive systems have been engineered in yeast (*13*), *D. melanogaster* (*7*), and two malaria vectors, *Anopheles stephensi* (*6*) and *An. gambiae* (*8*), with inheritance rates from heterozygotes reaching up to 99%.

Drive-resistant alleles are a significantly greater concern for homing-based systems than for *Medea*. For homing systems, resistant alleles emerge most commonly through error-prone copying and repair of the cleaved target site during the homing process. Of most concern are homing resistant alleles that confer no fitness cost to their host (*30*). These resistant alleles may spread at the expense of homing systems aimed at suppressing populations (*31*) and may also prevent homing systems aimed at driving desirable genes into populations from reaching fixation (*32*). Pre-existing genetic variation in a population is also a concern as, at a homing target site, a genetic variant could block the homing system from functioning (*30*). In fact, the CRISPR-Cas9-based homing systems engineered thus far have been compromised by resistant alleles in the first few generations of spread (*6, 8, 33, 34*), highlighting the need to develop drive architectures that prevent their rapid emergence (*1, 2, 31, 32*). Two methods proposed to achieve this are guide RNA multiplexing and targeting highly-conserved sites in essential genes (*1, 2, 31, 32*), and further work is required to thoroughly test these and other strategies.

### Driving Y chromosomes

In addition to the homing-based drive systems described above, CRISPR gene editing technology has advanced the potential for generating sex ratio-distorting driving Y chromosomes (*35*). These drives may function in XY species by expressing CRISPR components from Y chromosomes in the male germline to specifically cleave X chromosomes in many places during spermatogenesis (*28, 36*–*39*). This precisely-timed expression results in the destruction of X chromosomes during spermatogenesis, leading to an extreme male bias among offspring of males having the driving Y chromosome (Figure 1E). At the population level, an efficient and functional driving Y chromosome is expected to spread rapidly, eventually causing an all-male population crash. To date, CRIPSR-based Y-drive machinery has been demonstrated to function from an autosomal location in *An. gambiae*, and efforts are underway to transport it to the Y chromosome (*35*). Given that this system functions by cleaving many locations along the X chromosome, resistant chromosomes are less likely to evolve rapidly; however future research should investigate this more deeply and explore design features to further delay resistance evolution (*40*).

### Homing-based drive remediation systems

Before robust and efficient homing-based gene drive systems can be implemented in the wild, tools are required to remove the effector gene and possibly the entire drive system from the environment in the event of unwanted consequences. As for *Medea*, countermeasures have also been proposed for homing-based systems (*1, 2, 11*–*13*). These include an “ERACR” system (Element for the Reversal of the Autocatalytic Chain Reaction) consisting of a homing system with a target site corresponding to the original drive system, essentially removing the original drive system as it homes into it, and utilizing the Cas9 of the first system thus also removing this through the homing process (*12*) (Figure 2B). An “immunizing reversal drive” has also been proposed with its own Cas9 gene and target sites corresponding to both the original drive system and the wildtype allele (*1, 2*) (Figure 2C). This enables it to both remove the original drive system and to “immunize” wild-type individuals by homing into their corresponding target site, albeit while spreading another Cas9 gene into the population.

Additionally, a “CHACR” system (Construct Hitchhiking on the Autocatalytic Chain Reaction) has been proposed that utilizes the Cas9 from the first homing system to home into a second site in the genome in addition to the site of the original drive system, thus driving the CHACR into the second site while removing the original drive system, including its Cas9 gene, from the first site in the process (*12*) (Figure 2D). A fourth option is the release of individuals carrying an allele or chromosome conferring resistance to a homing or Y-drive system aimed at population suppression. This can remediate an environment of population-suppressing homing and Y-drive systems since those systems confer a large fitness load while resistance alleles do not and hence are selected for (*11, 31*).

### Population biology

Current work on countermeasures for homing systems is in its infancy, with molecular work just beginning and theoretical models only being analyzed in single, randomly mixing populations. Single-population models by Vella *et al.* highlight the oscillatory dynamics that can emerge for an ERACR system lacking its own Cas9 gene; however, the analysis of an immunizing reversal drive, and also second-generation *Medea* elements, requires consideration of population structure (*11*). A chasing dynamic in which the original homing or *Medea* system gets a lead on the immunizing or second-generation system is imaginable and could become apparent in a population consisting of multiple connected patches on a wide scale. Heterogeneity and chance are also essential considerations to model the dynamics of any threshold-independent drive remediation system. For instance, a few residual organisms having the original homing system could lead to it spreading back into the population once the majority wildtype state is restored. Structured population models such as those of North *et al.* and Eckhoff *et al.* could be used to explore these issues quantitatively, and further modeling will be required to gain insight into how these systems may behave in real populations (*41, 42*).

## Temporally self-limiting drive systems

### Killer-rescue

Another variety of drive system that could potentially be localized displays transient drive activity before being eliminated by virtue of a fitness cost. The simplest example of this is a two-locus system known as “killer-rescue” which consists of two alleles at unlinked loci - one that encodes a toxin (the killer allele) and another that confers immunity to the toxin (the rescue allele) (*15*). A release of individuals homozygous for both alleles results in temporary drive as the alleles segregate and the presence of the killer allele in the population confers a benefit to those also carrying the rescue allele. However, as the killer allele declines in frequency, the selective benefit of the rescue allele is lost and, if the rescue allele confers a fitness cost to the host, it is gradually eliminated from the population at a rate determined by the magnitude of its fitness cost. As discussed earlier, it is hoped that CRISPR technology will accelerate the development of these systems by providing new means to engineer toxins (*1*).

### Daisy drives

Daisy-chain drive is a variant of the CRISPR-based homing drive described earlier that has gained much attention recently due to its ability to function as a “local drive” (*2, 14, 32*). The system consists of a variable number of elements at unlinked sites arranged according to an architecture that results in temporally self-limiting drive. The first element has the effector gene (e.g. conferring disease refractoriness) and is driven by the Cas9 provided by the second element. The second element is driven by the Cas9 provided by the third element, and so on. The final element is not driven, and is expected to be eliminated from the population by virtue of its fitness cost. A release of individuals homozygous at all sites results in the effector gene being driven into the population; however the effect is transient as the final element falls out of the population and the others follow. The duration of drive depends on the number of daisy elements and their fitness costs. Drive resistance can be minimized by placing each element within an essential gene, expressing a recoded copy of that gene. A variant of this system is “daisyfield” drive, in which the first element is driven by Cas9 provided by a number of secondary elements, none of which are driven and all of which are eliminated from the population due to their fitness cost, also resulting in transient drive (*43*).

### Population biology

The dynamics of transient drive systems that rely on their fitness cost to be eliminated depends largely on the nature of the fitness cost. An optimal fitness cost may exist since there are competing demands - high fitness costs lead to rapid remediation following transient drive, but the maximum frequency that the system reaches in the population is compromised; while small fitness costs allow spread to higher maximum frequencies, but remediation from a population may take several years (*22*). Further complicating this, fitness costs are exceedingly difficult to quantify in the field, and even in the controlled environment of the laboratory, fitness costs have a tendency to change over time (*4, 5, 9*). It should also be noted that the spatial spread of temporally self-limiting drives is determined by how far the host species disperses while the drive system persists, and hence the degree of spatial dispersal is also determined by the fitness cost.

### Daisy quorum drive

Finally, it has recently been suggested that either daisy-chain or daisyfield drive systems could be used to induce underdominance in a population, thereby utilizing a temporally self-limiting drive system to create a system capable of threshold-dependent drive (*44*). The CRISPR gene editing that daisy drive systems rely upon can be used to insert genes at specific loci, and by using this ability to swap the locations of two haploinsufficient genes, underdominant dynamics could conceivably be induced. Once the daisy components of the system are lost, the underdominant system would be driven into populations in which it represents a majority and eliminated from populations in which it is a minority. This composite system, named “daisy quorum” drive, is yet to be engineered in the laboratory or to be rigorously explored even theoretically; but if it behaves as expected and the numerous components can be assembled in an evolutionarily stable way, it could be a valuable tool in the goal of local population replacement.

## Conclusions and future outlooks

In summary, the development of a wide range of gene drive systems will likely be accelerated through the application of CRISPR gene editing technology, encompassing a broad array of confinement properties. Threshold-dependent systems, such as chromosomal translocations and underdominant toxin-antidote systems, hold great promise for being limited to defined areas. Temporally self-limiting systems, such as killer-rescue and daisy-chain drives, hold promise for transient and hence local drive, although they are capable of diffusing into nearby populations and their eventual elimination is dependent on fitness costs, which are highly unpredictable in the wild. CRISPR technology has greatly accelerated the development of homing-based drives and driving Y chromosomes which are expected to be highly invasive once the evolution of resistance alleles is minimized. CRISPR has also accelerated the development of remediation systems for these drives; however, scenarios where homing systems outpace their countermeasures are easily imaginable, and limiting these systems might only be possible in highly contained environments.

Detailed ecological studies and population dynamic models, building upon those already developed, will be required to anticipate the confinement properties of these systems in wild populations. Lessons may be learned from recent releases of the intracellular bacterium *Wolbachia* as a biocontrol agent in *Ae. aegypti*. *Wolbachia* behaves analogously to a gene drive system, biasing inheritance in its favor through a mechanism known as cytoplasmic incompatibility, in which offspring of *Wolbachia*-infected mothers inherit the bacterium; but offspring of *Wolbachia*-infected fathers and uninfected mothers are unviable (*45*). This enables it to spread into a population from low initial frequencies; however it must exceed a low-to-moderate threshold frequency in order to drive in the presence of a fitness cost, which is often present. *Wolbachia* displayed patchy and incomplete spread through *Ae. aegypti* populations in two suburbs of Cairns, Australia (*25*); however it displayed rapid wave-like spread through *Drosophila simulans* populations in California (*46*). The difference is thought to be due to shorter dispersal and a higher fitness cost in *Ae. aegypti*, highlighting the relevance and need for species-specific data and models.

Future molecular and genetic work will be essential to continue developing the aforementioned CRISPR-based drive systems and to study their confinement properties, evolutionary stability and effectiveness at spread in contained laboratory populations. Given that each of the systems described here is susceptible to the evolution of drive resistance to varying degrees, it will be important to further investigate molecular designs that can impede the emergence of drive resistance. Through developing efficient and robust drive and remediation systems alongside an understanding of the ecosystems into which they are intended to spread, it is hoped that the many potential benefits of gene drive technology may one day be realized in a safe, ethical and responsible way.

## Acknowledgments

This work was supported by a private donation from www.MaxMind.com to OSA, US National Institutes of Health (NIH) K22 grant to OSA (5K22AI113060), a NIH R21 grant (1R21AI123937) to OSA, and a DARPA Safe Genes Program Grant (HR0011-17-2-0047) awarded to OSA and JMM.

## References

1. Champer, J., Buchman, A., and Akbari, O. S. (2016) Cheating evolution: engineering gene drives to manipulate the fate of wild populations, Nat. Rev. Genet. 17, 146–159.

2. Esvelt, K. M., Smidler, A. L., Catteruccia, F., and Church, G. M. (2014) Concerning RNA-guided gene drives for the alteration of wild populations, Elife 3.

3. Marshall, J. M., and Akbari, O. S. (2015) Gene drive strategies for population replacement.

4. Akbari, O. S., Matzen, K. D., Marshall, J. M., Huang, H., Ward, C. M., and Hay, B. A. (2013) A synthetic gene drive system for local, reversible modification and suppression of insect populations, Curr. Biol. 23, 671–677.

5. Buchman, A. B., Ivy, T., Marshall, J. M., Akbari, O., and Hay, B. A. (2016) Engineered reciprocal chromosome translocations drive high threshold, reversible population replacement in Drosophila.

6. Gantz, V. M., Jasinskiene, N., Tatarenkova, O., Fazekas, A., Macias, V. M., Bier, E., and James, A. A. (2015) Highly efficient Cas9-mediated gene drive for population modification of the malaria vector mosquito Anopheles stephensi, Proc. Natl. Acad. Sci. U. S. A. 112, E6736–43.

7. Gantz, V. M., and Bier, E. (2015) Genome editing. The mutagenic chain reaction: a method for converting heterozygous to homozygous mutations, Science 348, 442–444.

8. Hammond, A., Galizi, R., Kyrou, K., Simoni, A., Siniscalchi, C., Katsanos, D., Gribble, M., Baker, D., Marois, E., Russell, S., Burt, A., Windbichler, N., Crisanti, A., and Nolan, T. (2016) A CRISPR-Cas9 gene drive system targeting female reproduction in the malaria mosquito vector Anopheles gambiae, Nat. Biotechnol. 34, 78–83.

9. Akbari, O. S., Chen, C.-H., Marshall, J. M., Huang, H., Antoshechkin, I., and Hay, B. A. (2014) Novel synthetic Medea selfish genetic elements drive population replacement in Drosophila; a theoretical exploration of Medea-dependent population suppression, ACS Synth. Biol. 3, 915–928.

10. Chen, C.-H., Huang, H., Ward, C. M., Su, J. T., Schaeffer, L. V., Guo, M., and Hay, B. A. (2007) A synthetic maternal-effect selfish genetic element drives population replacement in Drosophila, Science 316, 597–600.

11. Vella, M. R., Gunning, C. E., Lloyd, A. L., and Gould, F. (2017) Evaluating Strategies For Reversing CRISPR-Cas9 Gene Drives.

12. Gantz, V. M., and Bier, E. (2016) The dawn of active genetics, Bioessays 38, 50–63.

13. DiCarlo, J. E., Chavez, A., Dietz, S. L., Esvelt, K. M., and Church, G. M. (2015) Safeguarding CRISPR-Cas9 gene drives in yeast, Nat. Biotechnol. 33, 1250–1255.

14. Noble, C., Min, J., Olejarz, J., Buchthal, J., Chavez, A., Smidler, A. L., DeBenedictis, E. A., Church, G. M., Nowak, M. A., and Esvelt, K. M. (2016) Daisy-chain gene drives for the alteration of local populations.

15. Gould, F., Huang, Y., Legros, M., and Lloyd, A. L. (2008) A killer-rescue system for self-limiting gene drive of anti-pathogen constructs, Proc. Biol. Sci. 275, 2823–2829.

16. Curtis, C. F. (1968) Possible use of translocations to fix desirable genes in insect pest populations, Nature 218, 368–369.

17. Laven, H., Cousserans, J., and Guille, G. (1972) Eradicating mosquitoes using translocations: a first field experiment, Nature 236, 456–457.

18. Lorimer, N., Hallinan, E., and Rai, K. S. (1972) Translocation homozygotes in the yellow fever mosquito, Aedes aegypti, J. Hered. 63, 158–166.

19. Vanoli, F., and Jasin, M. (2017) Generation of chromosomal translocations that lead to conditional fusion protein expression using CRISPR-Cas9 and homology-directed repair, Methods 121-122, 138–145.

20. Lekomtsev, S., Aligianni, S., Lapao, A., and Bürckstümmer, T. (2016) Efficient generation and reversion of chromosomal translocations using CRISPR/Cas technology, BMC Genomics 17.

21. Jiang, J., Zhang, L., Zhou, X., Chen, X., Huang, G., Li, F., Wang, R., Wu, N., Yan, Y., Tong, C., Srivastava, S., Wang, Y., Liu, H., and Ying, Q.-L. (2016) Induction of site-specific chromosomal translocations in embryonic stem cells by CRISPR/Cas9, Sci. Rep. 6, 21918.

22. Marshall, J. M., and Hay, B. A. (2012) Confinement of gene drive systems to local populations: a comparative analysis, J. Theor. Biol. 294, 153–171.

23. Davis, S., Bax, N., and Grewe, P. (2001) Engineered underdominance allows efficient and economical introgression of traits into pest populations, J. Theor. Biol. 212, 83–98.

24. Taylor, C., Touré, Y. T., Carnahan, J., Norris, D. E., Dolo, G., Traoré, S. F., Edillo, F. E., and Lanzaro, G. C. (2001) Gene flow among populations of the malaria vector, Anopheles gambiae, in Mali, West Africa, Genetics 157, 743–750.

25. Schmidt, T. L., Filipovic, I., Hoffmann, A. A., and Rasic, G. (2017) Fine-scale landscape genomics helps explain the slow spread of Wolbachia through the Aedes aegypti population in Cairns, Australia.

26. Beeman, R. W., and Friesen, K. S. (1999) Properties and natural occurrence of maternal-effect selfish genes (’Medea’ factors) in the red flour beetle, tribolium castaneum, Heredity 82 (Pt 5), 529–534.

27. Buchman, A., Marshall, J., Ostrovski, D., Yang, T., and Akbari, O. S. (2017) Synthetically Engineered Medea Gene Drive System in the Worldwide Crop Pest, D. suzukii.

28. Burt, A. (2003) Site-specific selfish genes as tools for the control and genetic engineering of natural populations, Proc. Biol. Sci. 270, 921–928.

29. Windbichler, N., Menichelli, M., Papathanos, P. A., Thyme, S. B., Li, H., Ulge, U. Y., Hovde, B. T., Baker, D., Monnat, R. J., Jr, Burt, A., and Crisanti, A. (2011) A synthetic homing endonuclease-based gene drive system in the human malaria mosquito, Nature 473, 212– 215.

30. Unckless, R. L., Clark, A. G., and Messer, P. W. (2017) Evolution of Resistance Against CRISPR/Cas9 Gene Drive, Genetics 205, 827–841.

31. Marshall, J. M., Buchman, A., Sánchez C, H. M., and Akbari, O. S. (2017) Overcoming evolved resistance to population-suppressing homing-based gene drives, Sci. Rep. 7, 3776.

32. Noble, C., Olejarz, J., Esvelt, K. M., Church, G. M., and Nowak, M. A. (2017) Evolutionary dynamics of CRISPR gene drives, Sci Adv 3, e1601964.

33. Hammond, A. M., Kyrou, K., Bruttini, M., North, A., Galizi, R., Karlsson, X., Carpi, F., D’Aurizio, R., Crisanti, A., and Nolan, T. (2017) The creation and selection of mutations resistant to a gene drive over multiple generations in the malaria mosquito.

34. Champer, J., Reeves, R., Oh, S. Y., Liu, C., Liu, J., Clark, A. G., and Messer, P. W. (2017) Novel CRISPR/Cas9 gene drive constructs reveal insights into mechanisms of resistance allele formation and drive efficiency in genetically diverse populations, PLoS Genet. 13, e1006796.

35. Galizi, R., Hammond, A., Kyrou, K., Taxiarchi, C., Bernardini, F., O’Loughlin, S. M., Papathanos, P.-A., Nolan, T., Windbichler, N., and Crisanti, A. (2016) A CRISPR-Cas9 sex-ratio distortion system for genetic control, Sci. Rep. 6, 31139.

36. Papathanos, P. A., Windbichler, N., and Akbari, O. S. Sex ratio manipulation for insect population control. In Transgenic insects: techniques and applications, pp 83–100.

37. Deredec, A., Burt, A., and Godfray, H. C. J. (2008) The population genetics of using homing endonuclease genes in vector and pest management, Genetics 179, 2013–2026.

38. Deredec, A., Godfray, H. C. J., and Burt, A. (2011) Requirements for effective malaria control with homing endonuclease genes, Proc. Natl. Acad. Sci. U. S. A. 108, E874–80.

39. Galizi, R., Doyle, L. A., Menichelli, M., Bernardini, F., Deredec, A., Burt, A., Stoddard, B. L., Windbichler, N., and Crisanti, A. (2014) A synthetic sex ratio distortion system for the control of the human malaria mosquito, Nat. Commun. 5, 3977.

40. Beaghton, A., Beaghton, P. J., and Burt, A. (2017) Vector control with driving Y chromosomes: modelling the evolution of resistance, Malar. J. 16, 286.

41. North, A., Burt, A., Godfray, H. C. J., and Buckley, Y. (2013) Modelling the spatial spread of a homing endonuclease gene in a mosquito population, J. Appl. Ecol. 50, 1216–1225.

42. Eckhoff, P. A., Wenger, E. A., Godfray, H. C. J., and Burt, A. (2017) Impact of mosquito gene drive on malaria elimination in a computational model with explicit spatial and temporal dynamics, Proc. Natl. Acad. Sci. U. S. A. 114, E255–E264.

43. Min, J., Noble, C., Najjar, D., and Esvelt, K. M. (2017) Daisyfield gene drive systems harness repeated genomic elements as a generational clock to limit spread.

44. Min, J., Noble, C., Najjar, D., and Esvelt, K. (2017) Daisy quorum drives for the genetic restoration of wild populations.

45. Werren, J. H., Baldo, L., and Clark, M. E. (2008) Wolbachia: master manipulators of invertebrate biology, Nat. Rev. Microbiol. 6, 741–751.

46. Turelli, M., and Hoffmann, A. A. (1991) Rapid spread of an inherited incompatibility factor in California Drosophila, Nature 353, 440–442.

